# Identification of candidate genes controlling black seed coat and pod tip color in cowpea (*Vigna unguiculata* L. Walp)

**DOI:** 10.1101/355586

**Authors:** Ira A Herniter, María Muñoz-Amatriaín, Sassoum Lo, Yi-Ning Guo, Timothy J Close

**Affiliations:** Department of Botany and Plant Sciences, University of California Riverside, Riverside, CA,USA

**Keywords:** Vigna unguiculata, MYB transcription factor, seed coat color, QTL analysis, SNP genotypinga

## Abstract

Seed coat color is an important part of consumer preferences for cowpea (*Vigna unguiculata* L. Walp). Color has been studied in numerous crop species and has often been linked to loci controlling the anthocyanin biosynthesis pathway. This study makes use of available resources, including mapping populations, a reference genome, and a high-density single nucleotide polymorphism genotyping platform, to map the black seed coat and purple pod tip color traits in cowpea. Several gene models encoding MYB domain protein 113 were identified as candidate genes. MYB domain proteins have been shown in other species to control expression of genes encoding enzymes for the final steps in the anthocyanin biosynthesis pathway. PCR analysis indicated that a presence/absence variation of one or more MYB113 genes may control the presence or absence of black pigment. A PCR marker has been developed for black seed coat color in cowpea.

## INTRODUCTION

Cowpea (*Vigna unguiculata* L. Walp) is a diploid (2n = 22) warm-season legume, mostly consumed as a grain, but also as a vegetable and often used as fodder for livestock. The seeds are used for cooking as whole beans or ground into a flour, while the immature pods and leaves are consumed as green vegetables (Singh 2014; Tijjani *et al*. 2015). Most cowpeas are grown by smallholder farmers under marginal conditions in sub-Saharan Africa, often as an intercrop (Ehlers and Hall 1997). In the United States, cowpeas are part of the traditional cuisine of the Southern states and are consumed as both fresh and dry beans (Fery 1985). Cowpea is a versatile crop due to its high adaptability to heat and drought, and its association with nitrogen fixing bacteria (Ehlers and Hall 1997). Over eight million tonnes were produced worldwide in 2013, with most of that production in Africa (FAOSTAT 2014, http://www.fao.org/faostat/en/#data/QC).

Seed coat color is an important consumer-related trait in cowpea. Previous research has indicated that consumers make decisions on the acceptability, quality, and presumed taste of a product depending on appearance, especially color (Kostyla *et al.* 1978; Simonne *et al.* 2001). Color preferences vary across and within markets as consumers prefer specific seed coat traits for different uses (Mishili *et al.* 2009). Newly developed varieties will be much more easily integrated into markets if the seeds are more visually similar to presently accepted varieties. As such, it behooves breeders to understand both the genetic basis of various seed coat traits and the consumer preferences in the markets to assist in breeding. Improved varieties often increase farmer income, which is frequently used for quality of life improvements, including education (Karanja *et al*. 2011).

Numerous genetic resources have been developed for use in cowpea. Among these are mapping populations including biparental recombinant inbred line (RIL) populations, an eight-parent Multiparent Advanced Generation InterCross (MAGIC) population, and a minicore population representing worldwide diversity of domesticated cowpea. Additionally, a genotyping array for 51,128 single nucleotide polymorphisms (SNP) was developed (Muñoz-Amatriaín *et al*. 2017) and a reference genome sequence of cowpea has been produced (Lonardi *et al.* in preparation, available in Phytozome [https://phytozome.jgi.doe.gov/]). Using these resources, consensus genetic maps of cowpea have been developed (Muchero *et al*. 2009; M. Lucas *et al.* 2011; Muñoz-Amatriaín *et al.* 2017) and major quantitative trait loci (QTL) for various traits have been mapped, including domestication-related traits (Lucas *et al*. 2015; Lo *et al*. 2018) and disease and pest resistance, among others.

Research on the inheritance of seed coat traits in cowpea began in the early 20^th^ century (Harland 1919; reviewed in Fery 1980). A factor called *Black seed color* (*Bl*) was identified through the study of F2 populations and found to also control sepal and pod tip color (Harland 1919, 1920). However, previous mapping efforts were hampered by the lack of high resolution mapping technologies and a reference genome. Here, we make use of these genetic and genomic resources to unveil the genetic basis of black seed coat and purple pod tip color and propose candidate genes.

## MATERIALS AND METHODS

### Plant Materials

Four populations were used for mapping: two biparental populations of RILs, an eight-parent Multiparent Advanced Generation InterCross (MAGIC) population (Huynh *et al.* 2018), and a minicore population representing the worldwide diversity of cultivated cowpea (Muñoz-Amatriaín *et al*. unpublished). One biparental population consists of 94 RILs developed at the University of California, Riverside, derived from a cross between “California Blackeye 27” (CB27), which has a medium-sized black eye seed coat and purple pod tips, and “IT82E-18,” which has a solid brown coat and green pod tips (Muchero *et al.* 2009). The other biparental RIL population was provided by the International Institute for Tropical Agriculture and consists of 121 RILs derived from a cross between “Sanzi,” a landrace with a speckled black and purple seed coat and purple pod tips, and “Vita 7,” which has a solid tan coat and green pod tips (Omo-Ikerodah *et al.* 2009). The seeds of each of these four parents are shown in Figure 1A. The MAGIC population consists of 305 RILs and was developed at the University of California, Riverside (Huynh *et al.* 2018). One of the eight parents of the population is CB27, which, as noted above, has a medium-sized black eye seed coat and purple pod tips. All three of the RIL populations segregate for black seed coat and purple pod tip color. The minicore population consists of 384 accessions and was developed at the University of California, Riverside (Muñoz-Amatriaín *et al*. unpublished). Accessions within the minicore population show great phenotypic diversity, including in seed coat color traits.

**Figure 1.**
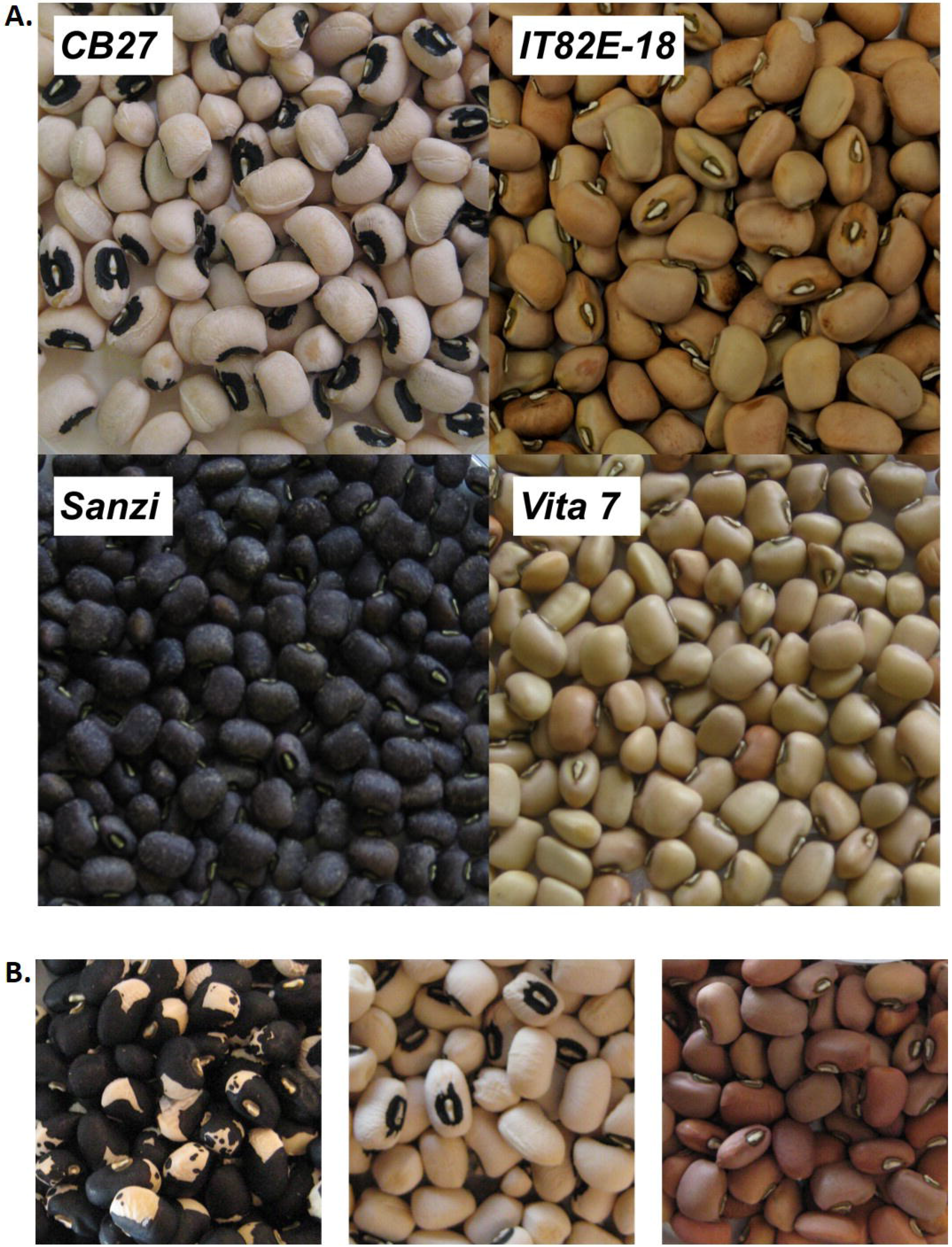
Seed coat color. (A) Images of the parents of the two biparental RIL populations. (B) A variety of seed coat color and patterns among the RILs from the CB27 by IT82E-18 population.

### SNP genotyping and data curation

DNA was extracted from young leaf tissue using the Qiagen DNeasy Plant Mini Kit (Qiagen, Germany) per the manufacturer’s instructions. The Cowpea iSelect Consortium Array (Illumina Inc., California, USA), which assays 51,128 SNPs (Muñoz-Amatriaín *et al.* 2017), was used to genotype each DNA sample. Genotyping was performed at the University of Southern California Molecular Genomics Core facility (Los Angeles, California, USA). The same custom cluster file as in Muñoz-Amatriaín *et al.* (2017) was used for SNP calling.

For the two biparental RIL populations, SNP data and genetic maps were available from Muñoz-Amatriaín *et al.* (2017). The CB27 by IT82E-18 genetic map included 16,566 polymorphic SNPs in 977 genetic bins, while the Sanzi by Vita 7 genetic map contained 15,619 SNPs in 1,275 genetic bins (Muñoz-Amatriaín *et al.* 2017). For the MAGIC population, SNP data and a genetic map were available from Huynh *et al.* (2018). The map included 32,130 SNPs in 1,568 genetic bins (Huynh *et al.* 2018). For the minicore population, a total of 41,514 SNPs were used after removing those with high levels of missing data and/or heterozygous calls (>20%), and with minor allele frequencies <0.05. SNPs in both the MAGIC and minicore populations were ordered based on their physical position in cowpea pseudomolecules (Lonardi *et al.* in preparation; [https://phytozome.jgi.doe.gov]).

### Phenotyping the populations

Phenotypic data for seed coat color were collected through visual examination of the seeds. Both biparental RIL populations and the MAGIC population segregated for black seed coat color. In the CB27 by IT82E-18 population lines were scored as “black” or “brown.” 21 lines were excluded due to missing seed coat data (Table S1). In the Sanzi by Vita 7 population lines were scored as “purple-black” or “tan” (Table S2). In both the MAGIC and the minicore populations lines were scored as “black” or “non-black” (Table S3, Table S4). Four lines in the MAGIC population were excluded due to missing seed coat data. Eleven accessions in the minicore that had no seed coat coloring were not included in the analysis as it is expected that this phenotype is due to a separate gene, known as *Color factor* (*C*) (Fery 1980). In all four populations black-seeded lines were given the score “1” while non-black-seeded lines were given the score “0.” Segregation distortion of the phenotypic data was assessed through chi-square tests. Pod tip color was examined through visual examination of immature seed pods in both biparental RIL populations and the MAGIC population; in every case, pod tip coloration was associated with black seed coat color.

### QTL and genome-wide association study (GWAS) analyses

QTL mapping in the biparental RIL populations was performed with the R packages “qtl” (Broman *et al.* 2003) and “snow” (Tierney *et al.* 2015). In “qtl” the function “read.cross” was used, which links the information from the phenotype and genotype files. Since the genetic map included many SNPs which mapped to the same cM position, the function “jittermap” was used, which randomly assigned each SNP a new map position by adding or subtracting a random value in the sixth decimal place. This enabled the use of all the SNP data in the QTL analysis. The probability value of each SNP was determined with the function “cal.genoprob(data, step=1),” from the “snow” package. Afterwards, to map the QTL probabilities, both a standard interval mapping using the EM algorithm: “scanone(data)” and a Haley-Knott regression: “scanone(data, method = “hk”, n.cluster=2)” were used. Both algorithms showed similar results. To test for significance, 1000 permutations were performed on the Haley-Knott regression: “scanone(data, method=“hk”, n.perm=1000).”

Marker effects were calculated first by using a hidden Markov model to simulate missing genotype data and to allow for genotyping errors: “sim.geno(cross = effectdata, n.draws = 16, step = 0, off.end = 0, error.prob = 0.001, map.function = “kosambi”, stepwidth = “fixed“)”. Then the effects were estimated across the genome: effectscan(cross = sim, get.se = FALSE)”. Percent variation explained by the identified QTL was determined by fitting the data to the putative QTL first by defining the QTL using the function “makeqtl(data, 5, 15.15, qtl.name = “bl”, what = “prob“)” (for the Sanzi by Vita 7 population 15.15 was replaced by 13.59), then using the function “fitqtl(data, qtl = bl, covar = NULL, method = “hk”, model = “binary“)”.

QTL mapping in the MAGIC population was performed using the R package “mpMap” (Huang and George 2011) with a protocol modified from that of Huynh *et al*. (2018). In short, the “mpIM” function was used with a step-length of 1 cM and a significance threshold of 8.096679e05, as determined through 1000 permutations of a null distribution: “mpIM(object=mp, ncov=0, responsename = trait, step=1, mrkpos=F, threshold=8.096679e-05, dwindow = 20” (Huynh *et al.* 2018). This function determined both the QTL probability and the effects from each parent as compared to one of the eight parents, IT93K-503-1.

GWAS was performed in the minicore population to identify SNPs associated with the black seed coat and purple pod tip color phenotype. The mixed-linear model (MLM) function (Zhang *et al.* 2010) implemented in TASSEL v.5 (https://www.maizegenetics.net/tassel) was used, with a principal component analysis (3 principal components) accounting for population structure in the dataset. The −log_10_(*p*) values were plotted against the physical coordinates of the SNPs, available from Phytozome (https://phytozome.jgi.doe.gov). A Bonferroni correction was applied to correct for multiple testing error in GWAS, with the significance cut-off set at α/n, where α is 0.05 and n is the number of tested markers (41,514). The marker effect was determined by taking the average of the MarkerR2 values of the significant SNPs multiplied by 100%.

### Candidate gene identification

Results from QTL and GWAS analyses were compared to identify the region containing overlap between significant regions in all four populations. The gene-annotated sequence of the overlapping QTL region was obtained from the reference genome sequence of cowpea (Lonardi *et al.* in preparation, [https://phytozome.jgi.doe.gov]). The list of genes in the overlapping region can be found in Table S5. Candidate genes were identified through similarity with genes responsible for similar traits in other species, including *Arabidopsis*, grape, citrus, and soybean, as determined by a review of the literature (see Discussion), as well as use of cowpea transcriptome data (Yao *et al.* 2016, [https://legumeinfo.org/]).

### PCR amplification

Primers were designed to amplify fragments at 5 kb and 1 kb intervals from the gene model *Vigun05g039500* to determine the size of the missing region in IT82E-18 (see Results). Further primers were designed to narrow the upstream and downstream edges of the deletion to ~1 kb and to amplify the MYB113 gene models affected by the deletion (see Results). All primer pairs, along with annealing temperatures, are listed in Table S6. PCR was performed using the Thermo Scientific DreamTaq Green PCR Master Mix (Thermo Scientific, Massachusetts, USA) per the manufacturer’s instructions. Primers were developed using Primer3 v0.4.1 (bioinfo.ut.ee/primer3) and ordered from Integrated DNA Technology (Coralville, Iowa, USA). PCR was run for 25-45 cycles with an annealing temperature compatible with the primer pair and an extension time of 60-75 seconds. PCR was performed on CB27 and IT82E-18 to determine the edges of the deleted region. Amplification to determine the presence or absence of affected MYB113 genes was performed on both a panel of lines from the CB27 by IT82E-18 population and in a set of ten diverse accessions with black and non-black seed colors from the minicore population (see Results). In the CB27 by IT82E-18 panel, the reference genome, IT97K-499-35, was used a positive control and water was used as a negative control. In the minicore panel IT97K-499-35-1 was used both as a positive control and as a representative of one of the major subpopulations identified by Muñoz-Amatriaín (unpublished). Amplicons were confirmed by gel electrophoresis.

### Data and Material Availability

Genotype data for the biparental RIL populations can be found in the supporting information Data S3 of Muñoz-Amatriaín *et al*. (2017). Genotype data for the MAGIC population can be found in the supporting information Data S1 of Huynh *et al*. (2018). Transcriptome data is available at https://legumeinfo.org. Genotype data for the minicore population is pending publication. Phenotype data for each population can be found in Table S1 (CB27 by IT82E-18), Table S2 (Sanzi by Vita 7), Table S3 (MAGIC), and Table S4 (minicore). The list of gene models in the shared significant region can be found in Table S5. Primer data can be found in Table S6. SNP LOD scores can be found in Table S7 (CB27 by IT82E-18) and Table S8 (Sanzi by Vita 7). cM LOD scores for the MAGIC population can be found in Table S9. SNP −log_10_(*p*) values for the minicore population can be found in Table S10. The overlapping SNPs with the highest significance can be found in Table S11 while the allele effects of the peak SNPs in the minicore can be found in Table S12. All supplemental tables have been uploaded to the GSA Figshare portal.

## RESULTS

### The genetic control of black seed coat and purple pod tip

In the CB27 by IT82E-18 population 47.3% (36) of tested lines had black seed coats while 52.7% (37) had brown seed coats (Table 1; examples in Figure 1B). In the Sanzi by Vita 7 population 57.0% (69) of tested lines had black seed coats while 43.0% (52) had tan colored seed coats (Table 1). In the MAGIC population 12.6% (38) of tested lines had black seed coats while 87.4% (263) had non-black seed coats. In the minicore population 28.2% (105) of tested accessions had black seed coats while 71.8% (268) had non-black colored seed coats (Table 1). Pod tip color was also scored in the CB27 by IT82E-18, Sanzi by Vita 7 and MAGIC populations, in all of which there was a perfect correlation with black seed coat color.

**Table 1.** SEED COAT COLOR PHENOTYPES FOR THE FOUR TESTED POPULATIONS. Included for each population are the number and percentage of lines with black seeds, the number with nonblack seeds, and those with no color or missing data.

| Population | # Black-seeded lines (% of tested lines) | # nonblack-seeded lines (% of tested lines) | # lines showing no color or missing data |
| --- | --- | --- | --- |
| *CB27* by *IT82E-18* | 36 (49.3%) | 37 (50.7%) | 0 |
| *Sanzi* by *Vita 7* | 69 (57.0%) | 52 (43.0%) | 0 |
| MAGIC | 38 (12.6%) | 263 (87.4%) | 4 |
| UCR minicore | 105 (28.2%) | 268 (71.8%) | 11 |

The biparental RIL populations and the MAGIC population demonstrated a seed coat color trait segregation not significantly different from the expected ratios of 1:1 in the biparental populations and 1:7 in the MAGIC population. The CB27 by IT82E-18 population had a chi-square value for a 1:1 ratio of 0.01 with a p-value of 0.92. The Sanzi by Vita 7 population had a chi-square value for a 1:1 ratio of 2.39 with a p-value of 0.12. The MAGIC population had a chi square value for 1:7 of 0.004 with a p-value of 0.95. The near 1:1 segregation in the biparental RIL populations and the near 1:7 segregation in the MAGIC population indicate that there is likely a single region which controls black seed coat color and purple pod tip color in the populations, consistent with the findings of Harland (1919, 1920).

### Black seed coat and purple pod tip mapping

Following phenotypic characterization of the seed coat color, QTL were identified using the R package “qtl” (Broman 2009) for the biparental RIL populations, the R package “mpMap” (Huang and George 2011) for the MAGIC population, and the MLM method in TASSEL v5 (maizegenetics.net/tassel) for the minicore (see Methods for more details). These methods determined a QTL interval of 30.92 cM (corresponding to 8,689,246 bp in the cowpea reference sequence) in the CB27 by IT82-18 population (Figure 2A), 39.23 cM (16,358,257 bp) in the Sanzi by Vita 7 population (Figure 2B), an interval of 2 cM (607,087 bp) in the MAGIC population (Figure 2C), and 1,087,245 bp in the minicore population (corresponding to 1.81 cM in the consensus map by Muñoz-Amatriaín, *et al*. [2017] [Figure 2D]). Significant regions and flanking markers can be found in Table 2. All four QTL mapped to the same region on Vu05, allowing a narrowing of the QTL region to the size of the region contained within all four QTL, between the SNPs 2_12036 and 2_15997, a range of 273,283 bp. The percent variation explained by the QTL in both biparental RIL populations was 75.00%. The QTL effect was 0.50 in the CB27 by IT82-18 population and 0.48 in the Sanzi by Vita 7 population. In the MAGIC population, the QTL explained 72.19% of the variation, with most of the effect coming from the black parent (0.93, CB27). In the minicore population, the QTL explained was 14.61% of the variation. The overlapping SNPs with the highest significance can be found in Table S11 while the allele effects of the peak SNPs in the minicore can be found in Table S12.

**Figure 2.**
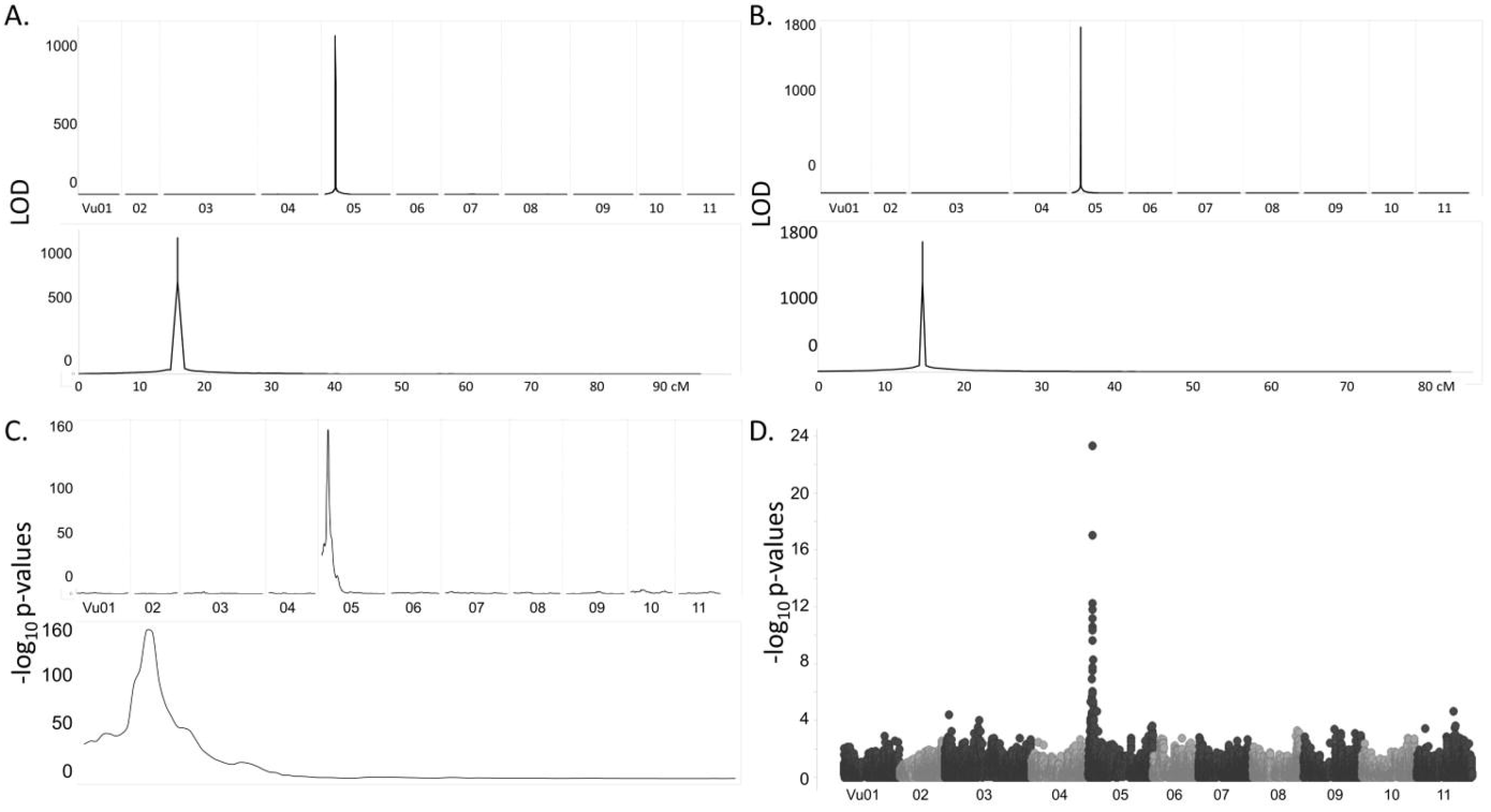
Mapping of the black seed coat trait. (A) QTL mapping in the CB27 by IT82E-18 population. (B) QTL mapping in the Sanzi by Vita 7 population. (C) GWAS analysis of the minicore population.

**Table 2.**
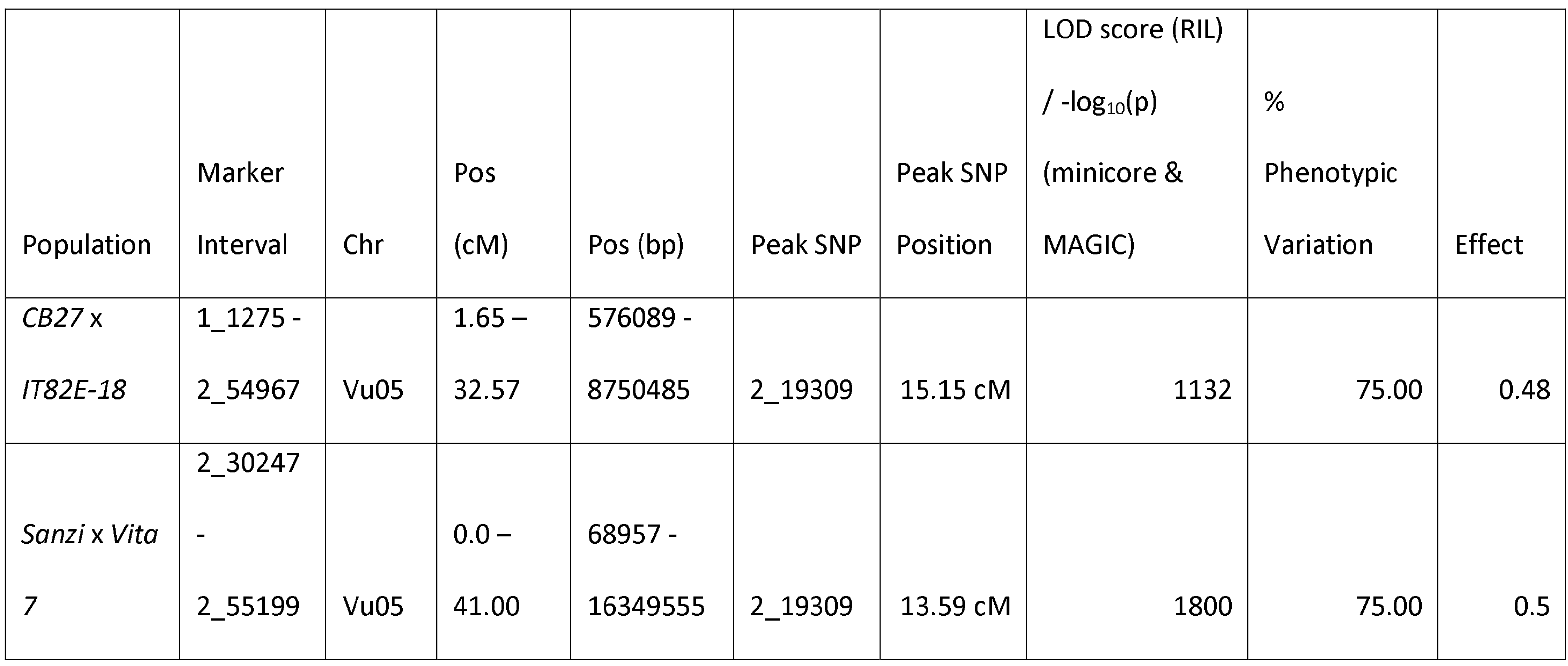
SIGNIFICANT QTL IDENTIFIED IN THE RIL AND MINICORE POPULATIONS. For each population the marker interval of significant SNPs (LOD>3.22 in the RIL populations, −log10(*p*)>5.92 in the minicore population), the chromosome the QTL on which the QTL is located, the peak SNP, the position of the peak SNP (on the genetic map used for the RIL populations and on the physical map for the minicore population), the score of the peak SNP (LOD in the RIL populations, −log10(*p*) in the minicore, the phenotypic variation explained by the QTL, and the QTL effects are shown.

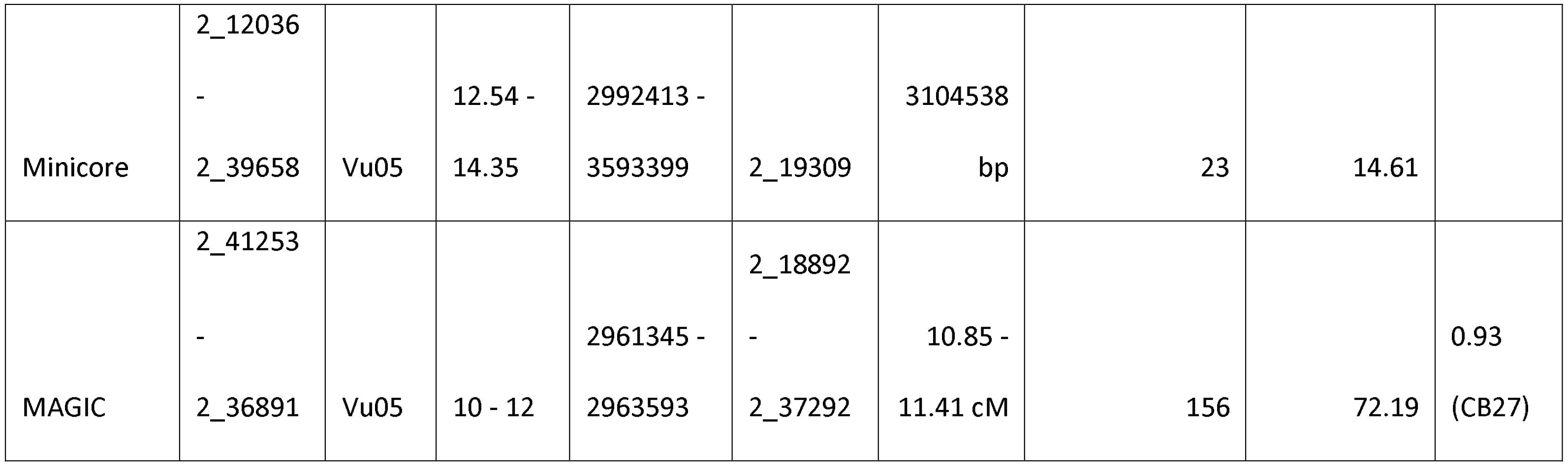

### Identification of candidate genes

The overlapping QTL region of 273,283 bp contains thirty-five gene models in the reference genome (Table S5). Upon further examination of the peak region in the minicore population, it was noted that between the two SNPs with the highest −log_10_(*p*) values (2_19309 and 2_15182) there are only thirteen gene models. Among the thirteen gene models are five coding for MYB domain protein 113, hereafter referred to as “MYB113”: *Vigun05g039300*, *Vigun05g039400*, *Vigun05g039500*, *Vigun05g039700*, and *Vigun05g039800*. Based on previous studies, the MYB gene family has been identified as being a regulator of genes involved in the anthocyanin biosynthesis pathway and in pigmentation in a wide range of other plants (see Discussion), and so the MYB113 genes were considered strong candidates. The expression profiles of the five gene models were examined at the Legume Information System portal (http://legumeinfo.org), using data from Yao *et al.* (2016) (Figure S1). Of the five, *Vigun05g039400* and *Vigun05g039500* showed high expression levels in the developing seeds, with *Vigun05g039500* showing much higher expression than *Vigun05g039400*, *Vigun05g039300* showed high expression in the developing pods, flowers, and in leaves, while Vigun*05g039800* showed high expression in the leaves and lower expression in the stem. *Vigun05g039700* showed no expression in any of those tissues. Between *Vigun05g039500* and *Vigun05g039700* is another gene model, *Vigun05g039600*. However, that gene model encodes an EXS family protein, which is mostly expressed in root tissue. There is no prior literature associating such a gene with pigmentation, and so it was not considered to be a candidate gene. The expression data suggest that *Vigun05g039400* and *Vigun05g039500* control the black seed coat color, while *Vigun05g039300* controls the purple pod tip color.

### Amplification of candidate genes

Due to its high expression level in the seed (Figure S1), *Vigun05g039500* was the first candidate gene chosen for further analysis by amplification and sequencing to search for allelic differences. PCR was performed on segments of *Vigun05g039500* in a panel of parents and lines from the CB27 by IT82E-18 population. The results showed a consistent successful amplification in all black-seeded lines and a similarly consistent failure to amplify in brown-seeded lines, indicating a possible presence/absence variation (Figure 3C).

**Figure 3.**
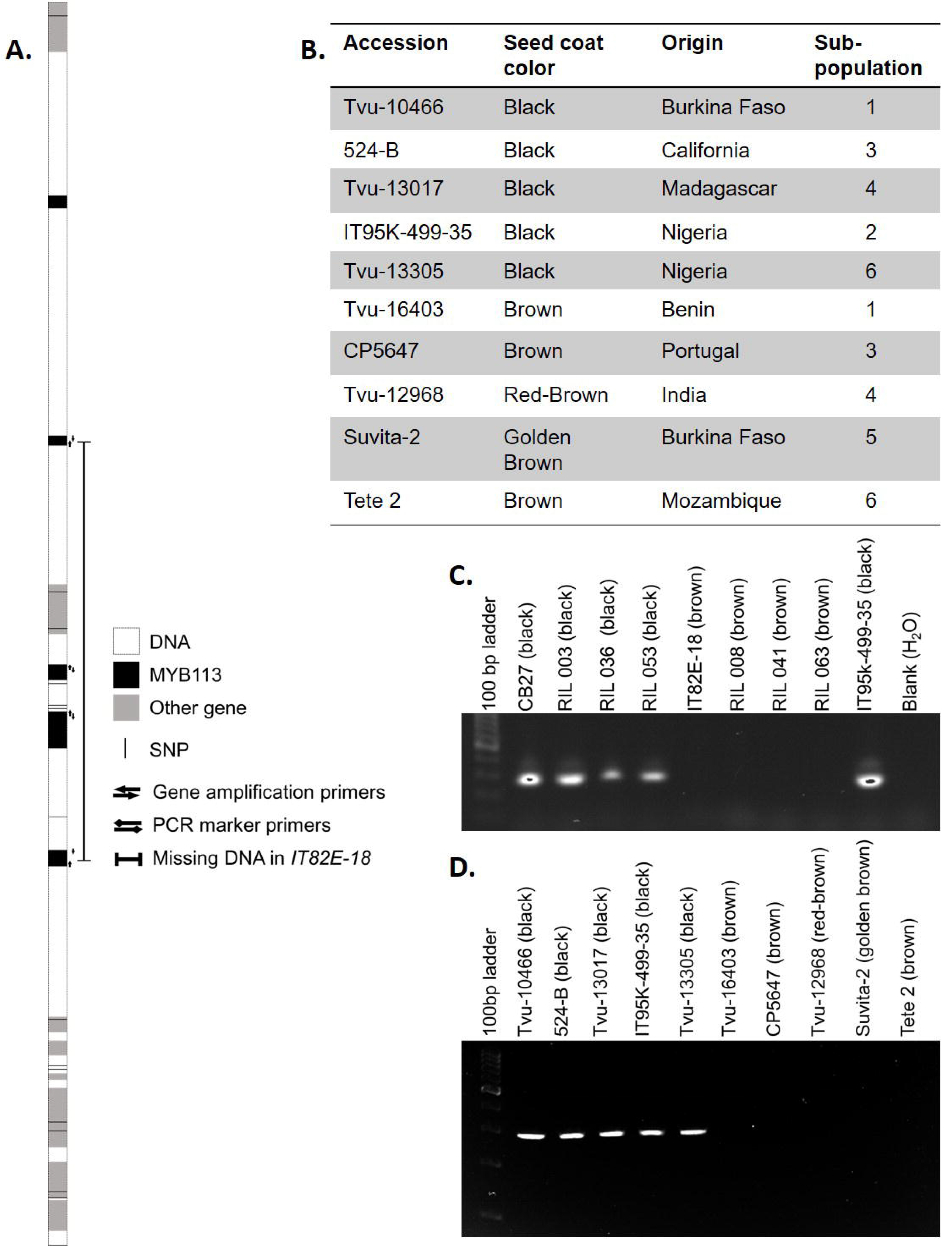
Gene identification. (A) Diagram of the peak significance region, including SNPs from the iSelect SNP genotyping platform (black lines), genes encoding MYB113 (black boxes), other local gene models (grey box) and the extent of the deletion in the IT82E-18 (bar with block ends). Other notations are indicated in the figure. (B) Information of minicore accessions used for validation. C) PCR results from the marker primers designed to amplify a 278 bp segment in the largest exon of *Vigun05g039500* in the CB27 by IT82E-18 population (top) and the minicore panel (bottom).

A list of nearly one million SNPs developed for the purpose of designing the Cowpea iSelect genotyping platform (Muñoz-Amatriaín *et al*. 2017) was examined for evidence of a possible presence/absence variation consisting of a deletion between black and non-black seeded lines (Figure 4). The SNP list showed a clear distinction between the two groups, with a block of failed SNPs in most of the non-black seeded lines extending from 3,137,965 bp to 3,176,886 bp in chromosome Vu05, supporting a presence/absence variant. There was one exception to the pattern, 24-125-B-1, which has a brown eye, but has successful SNP calls in the missing section. PCR was performed on small DNA segments about every 5 kb in both directions from *Vigun05g039500* in both CB27 and IT82E-18 until amplification was successful in both lines.

**Figure 4.**
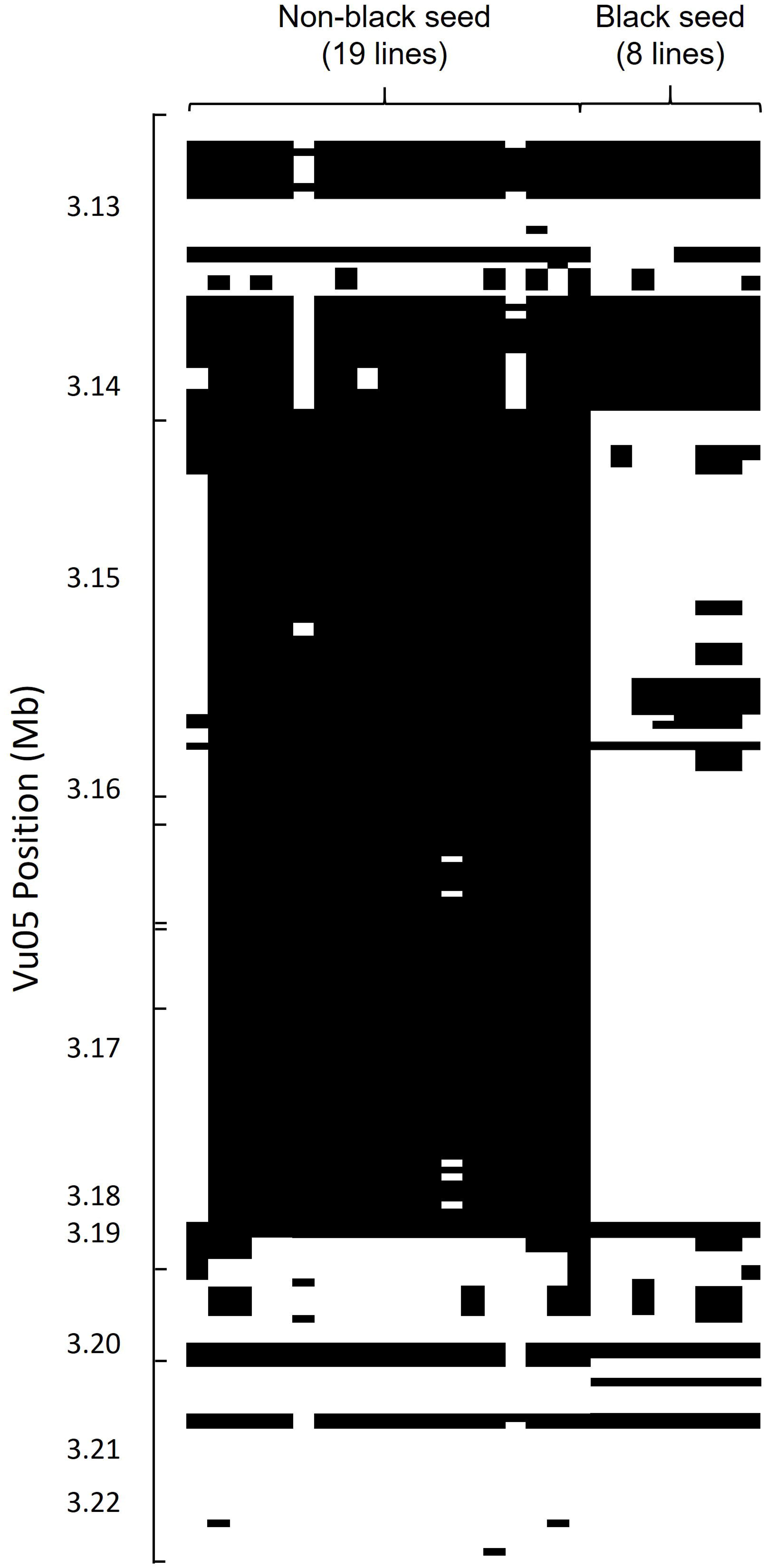
Assessing the deleted area with data from the SNP discovery panel. SNPs that were identified from 37 diverse accessions (Muñoz-Amatriaín *et al.* 2017) are arranged by physical position with markers included in the iSelect array noted. Accessions are arranged based on seed coat color. Absence of the DNA sequence in the SNP position is indicated by black color. Black areas therefore represent missing DNA sequence regions. Tic marks indicate SNP markers included in the iSelect Cowpea Consortium Array.

Then PCR was performed in about 1 kb intervals to further narrow the edges of the deletion, and then in smaller intervals to determine the edges more precisely. It was determined by doing so that a segment of about 40 to 42 kb in length, beginning between 3,142,209 and 3,143,232 bp (1,023 bp range) and ending between 3,183,152 and 3,184,076 bp (924 bp range) on Vu05, is present in the reference genome and in CB27 and is absent in IT82E-18 (Figure 3A). This puts the edges of the missing region inside the genomic sequences of *Vigun05g039300* and *Vigun05g039700*, indicating that three genes are missing entirely (two MYB113 genes, *Vigun05g039400* and *Vigun05g039500*, and an EXS gene, *Vigun05g039600*) and two are truncated (two MYB113 genes *Vigun05g039300* and *Vigun05g039700*) in IT82E-18 (Figure 3A).

The five MYB113 gene models in the cluster were compared via BLAST to one another to determine levels of similarity. The e-scores of the pairwise comparisons ranged from 0.0 to 3.00e^−69^. Based on the results, *Vigun05g039300* and*Vigun05g039700*, are the most similar (e-score = 0.0, 97% identity), followed by *Vigun05g039300* and *Vigun05g039800* (e-score = 0.0, 96% identity)

### Validation of candidate genes

To clarify which MYB113 gene/s might be required for the expression of black seed coat and purple pod-tip color, PCR amplification was performed on two panels, one of lines from the CB27 by IT82E-18 population and one of diverse accessions from the minicore, using primers developed to uniquely amplify each MYB113 gene affected by the deletion. The CB27 by IT82E-18 panel consisted of the parents and three lines each with black or brown seeds. The minicore panel consisted of ten lines, two each from each of the major subpopulations identified by Muñoz-Amatriaín (unpublished), with one black-seeded line and one non-black-seeded line from each (Figure 3B). Included as a positive control in both panels as well as a representative of one of the subpopulations in the minicore panel was the cowpea reference genome, IT97K-499-35. Whole gene amplification was performed in *Vigun05g039300* and *Vigun05g039700*, while segments of the largest exon were amplified in each of both *Vigun05g039400* and *Vigun05g039500*. Amplification of all four tested MYB113 genes succeeded in all black-seeded lines of the CB27 by IT82E-18 panel and failed in all brown seeded lines (Figure 3C, Figure S2). Amplification of *Vigun05g039300* and *Vigun05g039400* was successful in only two of the five tested black-seeded accessions, indicating that the presence of either of these genes is not required for black seed coat color. Amplification of *Vigun05g039500* was successful in all black-seeded accessions, as was amplification of *Vigun05g039700*. Amplification failed for all tested primer pairs in all non-black-seeded accessions in the minicore panel (Figure 3D, Figure S2). The inconsistent amplification of
*Vigun05g039300* and *Vigun05g039400* indicates possible variability in the size of the deleted region.

## DISCUSSION

Anthocyanins are plant pigments which are produced in numerous plant organs, including flowers, fruits, and seeds and are known to be a major source of coloring in seed coats, with different molecules known to be responsible for various colors (Petroni and Tonelli 2011). The candidate genes identified in this analysis, the MYB113 genes on chromosome Vu05, belong to the R2R3 MYB class of transcription factors. The MYB transcription factor family, and especially the R2R3-MYB subfamily has been implicated in plant pigment production in various tissues (Liu *et al.* 2015). R2R3 MYBs, so named for their two MYB DNA-binding domains, function as part of a modular complex in conjunction with a helix-loop-helix protein and a WD repeat protein (Liu *et al.* 2015). This modular function, and especially the interchangeability of the R2R3 MYBs is consistent with observations in *Arabidopsis* (Liu *et al.* 2015), grape (Kobayashi 2004; Walker *et al.* 2007) and citrus (Butelli *et al*. 2012). Proteins which have been shown to be regulators of genes involved in the anthocyanin biosynthesis pathways include the products of *Arabidopsis* genes *AT1G66370*, *AT1G66380*, and *AT1G66390* (Liu *et al*. 2015), grape genes *VvMYBA1* and *VvMYBA2* (Walker *et al*. 2007), and the soybean gene *Glyma.09G235100* (Yan *et al.* 2015). These genes are homologous to the MYB113 genes and, similar to the cowpea MYB genes, the genes in other systems are clustered together, lending further credence to the similarity between systems. Interruption of these R2R3-MYBs, often caused by an transposable element insertion, can result in a change in the observed color, as in grape (Kobayashi 2004; Walker *et al*. 2007), citrus (Butelli *et al*. 2012), and soybean (Yan *et al*. 2015).

The expression data of the MYB113 genes showed that *Vigun05g039400* and *Vigun05g039500* were relatively highly expressed in developing seeds while Vigun05g039300 was most highly expressed in the pods, flowers, leaves. Additionally, the inconsistent presence of the Vigun05g039300 and Vigun05g039400 in the minicore panel (Figure S2) indicates that the presence of either of these genes is not required for black seed coat color. It is possible that when either *Vigun05g039400* or *Vigun05g039500* is involved in the complex it causes upregulation of genes encoding enzymes in the anthocyanin biosynthesis pathway in the seed coat while when *Vigun05g039300* is involved upregulation of the pathways in the seed pod tip. The physical closeness of the genes would explain the observed perfect correlation between black seed coat and purple pod tip coloring. Further research on the MYB113 genes is needed to confirm the genes’ roles in seed coat and pod tip color through transient or stable expression in lines that normally do not express the pigmentation.

In the present analysis it was determined that the deletion in IT82E-18 begins between 3,142,209 and 3,143,232 bp and ends between 3,183,152 and 3,184,076 bp on Vu05. Wild cowpea accessions tend to have black seed coat color. This suggests that IT82E-18 carries an abnormal mutation, in this case a deletion, which may have been selected for by cultivators who noticed unusual seed colors. BLAST results comparing the genomic sequence of the MYB genes indicate that *Vigun05g039300*, *Vigun05g039700*, and *Vigun05g039800* are highly similar, with *Vigun05g039300* and *Vigun05g039700* the most similar among the five gene models. The deletion appears to begin in *Vigun05g039300* and end in *Vigun05g039700*, and it could have arisen through non-allelic homologous recombination, an unequal crossover between highly similar DNA sequences, as described by Gu *et al.* (2008). Other accessions may have a different number of MYB113 genes than the sequenced reference genome, IT97K-499-35 (Lonardi *et al.* in preparation [https://phytozome.jgi.doe.gov/]). Similarly, it is possible that in other accessions, the size of the deletion may vary.

One of the lines used for determining the size of the missing region from the SNP design panel, 24-125-B-1, has a small brown eye. However, unlike the rest of the non-black seeds in the panel, it has alignment data in the region missing in the other accessions (Figure 4). This may indicate that the lack of black pigmentation in the line is caused by a different variation. Further experimentation is needed to determine the genetic basis of the lack of pigmentation in the line.

A recently assembled reference genome was used to determine the sequence of the MYB113 gene models (Lonardi *et al.* in preparation [https://phytozome.jgi.doe.gov/]). This reference genome was assembled using DNA from the variety IT97K-499-35, which is a black-eye seeded variety. Had the reference genome sequence been developed from a variety with a non-black seed coat, such as IT82E-18 or Vita 7, it would have been more complicated to identify the candidate gene, to design the primers used to determine the size of the deletion, or to develop as a PCR marker for black seed coat color. Additionally, the list of nearly one million SNPs that were identified during development of the Cowpea iSelect Consortium Array (Muñoz-Amatriaín *et al*. 2017) was instrumental in determining the edges of the deleted region, as well as to show that the deletion is widespread among cultivated cowpeas. Current efforts to gain insights into the cowpea pan-genome by sequencing additional accessions could shed further light on the variation of this region in cultivated cowpea.

## CONCLUSIONS

Advances in genomics over the past several years have enabled the elucidation the genetic basis of black seed coat and purple pod tip color, traits first described nearly one hundred years ago
(Harland 1919, 1920). This study maps black seed coat and purple pod tip color in several independently generated populations and provides candidate genes. Using high-throughput SNP genotyping and whole genome sequencing, previously impossible levels of mapping precision have been achieved. The presence of a deletion is supported by PCR evidence and sequence alignment from thirty-seven accessions. The identification of MYB transcription factors as candidate genes is supported by prior literature on homologous genes performing similar functions in other species, including *Arabidopsis*, grape, citrus, and soybean. The PCR-based markers developed here provide a useful tool for breeders engaged in marker-assisted selection for seed coat color in cowpea.

## ACKNOWLEDGEMENTS

Authors thank: Alex Rajewski for developing a tutorial for QTL mapping with R/qtl; Mitchell Lucas and William Moore for preliminary genetic mapping of the black color gene in the Sanzi by Vita 7 RIL population; Christian Fatokun for providing the Sanzi by Vita 7 population; Zhenyu Jia for assistance with quantitative analysis; Steve Wanamaker for technical support and assistance; Bao-Lam Huynh for assistance with mapping in the MAGIC population; and the University of Southern California Genomic Core team for iSelect genotyping services. This study was supported by the Feed the Future Innovation Lab for Climate Resilient Cowpea (USAID Cooperative Agreement AID-OAA-A-13-00070), the National Science Foundation BREAD program (Award #1543963), and Hatch Project CA-R-BPS-5306-H.

